# Quantitative cancer-immunity cycle modeling for predicting disease progression in advanced metastatic colorectal cancer

**DOI:** 10.1101/2024.04.30.591845

**Authors:** Chenghang Li, Yongchang Wei, Jinzhi Lei

**Affiliations:** School of Mathematical Sciences, Tiangong University, Tianjin, China; Department of Radiation and Medical Oncology, Zhongnan Hospital of Wuhan University, Wuhan University, Wuhan, China; Hubei Key Laboratory of Tumor Biological Behaviors, Zhongnan Hospital of Wuhan University, Wuhan University, Wuhan, China; Center for Applied Mathematics, Tiangong University, Tianjin, China

## Abstract

Patients diagnosed with advanced metastatic colorectal cancer (mCRC) often exhibit heterogeneous disease progression and face poor survival prospects. In order to comprehensively analyze the varied treatment responses among individuals and the challenge of tumor recurrence resistant to drugs in advanced mRCR, we developed a novel quantitative cancer-immunity cycle (QCIC) model. The proposed model was meticulously crafted utilizing a blend of differential equations and randomized modeling techniques to quantitatively elucidate the intricate mechanisms governing the cancer-immunity cycle and forecast tumor dynamics under different treatment modalities. Furthermore, by integrating diverse clinical datasets and rigorous model analyses, we introduced two pivotal concepts: the treatment response index (TRI) and the death probability function (DPF). These concepts are crucial tools for translating model predictions into clinically relevant evaluation indexes. Using virtual patient technology, we extrapolated tumor predictive biomarkers from the model to predict survival outcomes for mCRC patients. Our findings underscore the significance of tumor-infiltrating CD8+CTL cell density as a key predictive biomarker for short-term treatment responses in advanced mCRC while emphasizing the potential predict value of the tumor-infiltrating CD4+Th1/Treg ratio in determining patient survival. This study presents a pioneering methodology bridging the divide between diverse clinical data sources and the generation of virtual patients, offering invaluable insights into understanding inter-individual treatment variations and forecasting survival outcomes in mCRC patients.

**Author summary:** This study introduces a sophisticated modeling approach to delineate the intricate dynamics of tumor-immune interactions within the cancer-immunity cycle. Utilizing a multi-compartmental, multi-scale, multi-dimensional differential equation model, we quantified the complex interplay between tumor cells, immune cells, cytokines, and chemokines. By integrating virtual patient technology, we have conducted *in silico* clinical trials that accurately predict disease progression across multiple treatment modalities for cancer patients, particularly in advanced mCRC. Through the combination of differential equations and randomized modeling, we effectively captured the diverse treatment responses and clinical manifestations of drug-resistant tumor recurrence. Furthermore, we explored the pivotal role of predictive biomarkers in determining patients’ survival prognosis, offering insights for tailored therapeutic strategies. Importantly, our computational framework holds promise for the extension to the investigation of patients with other tumor types, thus contributing to the personalized investigation of patients with other tumor types and cancer care.

## Introduction

Colorectal cancer (CRC) ranks as the third most prevalent malignant tumor globally and the second leading cause of cancer-related mortality, representing a significant health challenge [1]. Metastatic colorectal cancer (mCRC) is particularly associated with poor prognosis, posing considerable clinical hurdles [2, 3]. Recent clinical trials have highlighted the efficacy of combination therapy involving the novel chemotherapeutic TAS-102 and bevacizumab in advanced mCRC patients, substantially extending survival rates [4]. This therapeutic approach is expected to revolutionize the treatment of advanced mCRC based on its promising results. However, despite the favorable efficacy demonstrated by TAS-102 plus bevacizumab combination therapy compared to standard treatments, efficacy discrepancy and tumor recurrence remain the major challenges in clinical treatment [5].

The pivotal role of tumor-immune interactions in cancer initiation and progression underscores the importance of understanding these dynamics [6–8]. The recently proposed concept of the cancer-immunity cycle offers a comprehensive framework that elucidates the pivotal role of the immune system in eradicating tumor cells [9–11].

Unlike the previous focus solely on tumor-immune interactions within the tumor microenvironment (TME), the cancer-immunity cycle comprehensively outlines the immune response process across various body compartments [9–11]. Understanding the quantitative dynamics of the seven steps of the cancer-immunity cycle is crucial for the prognostic prediction in mCRC patients undergoing combination therapy.

Mathematical modeling offers a powerful approach for elucidating the complex interactions between tumors and immunity. In the 1990s, Kuznetsov et al. [12] applied the foundational principles of the Lotka-Volterra model to the tumor-immunity paradigm. Subsequently, Kirschner et al. [13] integrated the interactions between IL-2, tumor cells, and effector cells. Pillis et al. [14–16] introduced NK cells, CD8+T cells, and circulating lymphocytes into their modeling framework, further exploring the impact of chemotherapy and immunotherapy on tumor evolution. Robertson-Tessi et al. [17, 18] proposed a comprehensive mathematical model of tumor-immunity interactions, incorporating a negative feedback loop to account for immunosuppressive mechanisms. Additionally, numerous studies have developed mathematical models to investigate various aspects of tumor-immunity interactions, thus expanding the breadth of research in this field [19–28]. While attempting to encompass all the cell types and signaling molecules involved in tumor-immunity interactions may be ambitious, it is crucial to strike a balance, as overly simplistic models may fail to capture the complex dynamics observed both experimentally and clinically.

In recent years, significant advancements in tumor immunology have led to the establishment of high-dimensional complex models based on the mechanism of tumor immunity. Lai and Friedman et al. [29–37] elucidated the biological mechanisms underlying interactions among tumor cells, immune cells, and cytokines through reaction-diffusion, facilitating the development of mathematical models for combination therapy in tumors. These studies primarily aimed to investigate the synergistic or antagonistic effects of different drugs in tumor therapy using mathematical frameworks. Notably, they demonstrated that non-overlapping dosing regimens yield significantly greater benefits in combination therapy compared to concurrent dosing methods. Additionally, these models explored the impact of different dosing sequences on tumor therapy, offering insights applicable to various cancer types and aiding in the design of clinical trials. However, these models did not delve into inter-individual treatment differences among patients.

Conversely, significant attention has been directed toward quantitative systems pharmacology (QSP) models, which elucidate tumor-immunity interactions using ordinary differential equations (ODE) and simulate complex inter-cellular regulation through agent-based models (ABM) [38–50]. These models compartmentalize the human body into four regions: central, peripheral, tumor, and tumor-draining lymph nodes, incorporating diverse immune cells, cytokines, and drug modules into respective compartments to emulate real immune responses. These investigations aim to elucidate inter-individual treatment disparities among patients receiving identical therapies.

Moreover, established QSP models have demonstrated the capacity to identify predictive biomarkers for tumors, crucial for assessing short-term treatment outcomes based on biomarker expression. However, while these models focus on elucidating drug mechanisms of action, they do not provide a comprehensive quantitative depiction of the cancer-immunity cycle.

In this study, we present a quantitative cancer-immunity cycle (QCIC) model, a multi-compartmental, multi-scale, multi-dimensional ODE model, to elucidate the complex dynamics of tumor-immune interactions during disease progression and therapy. Integrating knowledge from immunology, oncology, clinical medicine, computational systems biology, and applied mathematics, the QCIC model provides a detailed mathematical description of cell fate decisions and tumor immunity mechanisms. The QCIC model provides insight into personalized therapeutic strategies by accurately capturing inter-individual treatment variation. Additionally, the QCIC model facilitates accurate efficacy predictions for population-level clinical trials through virtual patient technology [51–54]. We introduce the treatment response index (TRI) to quantify disease progression in virtual clinical trials and the death probability function (DPF) to predict overall survival (OS). Furthermore, we explore the impact of predictive biomarkers on survival prognosis in advanced mCRC patients, identifying tumor-infiltrating CD8+CTL cells and the tumor-infiltrating CD4+Th1/Treg ratio as significant predictors of disease progression and survival prognosis, respectively.

## Results

### Research framework and the quantitative cancer-immunity cycle model

In this study, we developed a mathematical model based on the cancer-immunity cycle theory, termed the quantitative cancer-immunity cycle (QCIC) model, to quantitatively describe the dynamic evolution of metastatic colorectal cancer (mCRC) [9–11]. The establishment of the QCIC model facilitates the following objectives: 1) capturing the phenomenon of drug resistance and recurrence in mCRC patients during clinical treatment, 2) understanding the inter-individual therapeutic disparities among mCRC patients receiving identical treatment regimens, and 3) predicting the survival rate of mCRC patients using predictive biomarkers extracted from the QCIC model. A schematic of the research framework and the QCIC model is presented in Figure 1.

**Fig 1.**
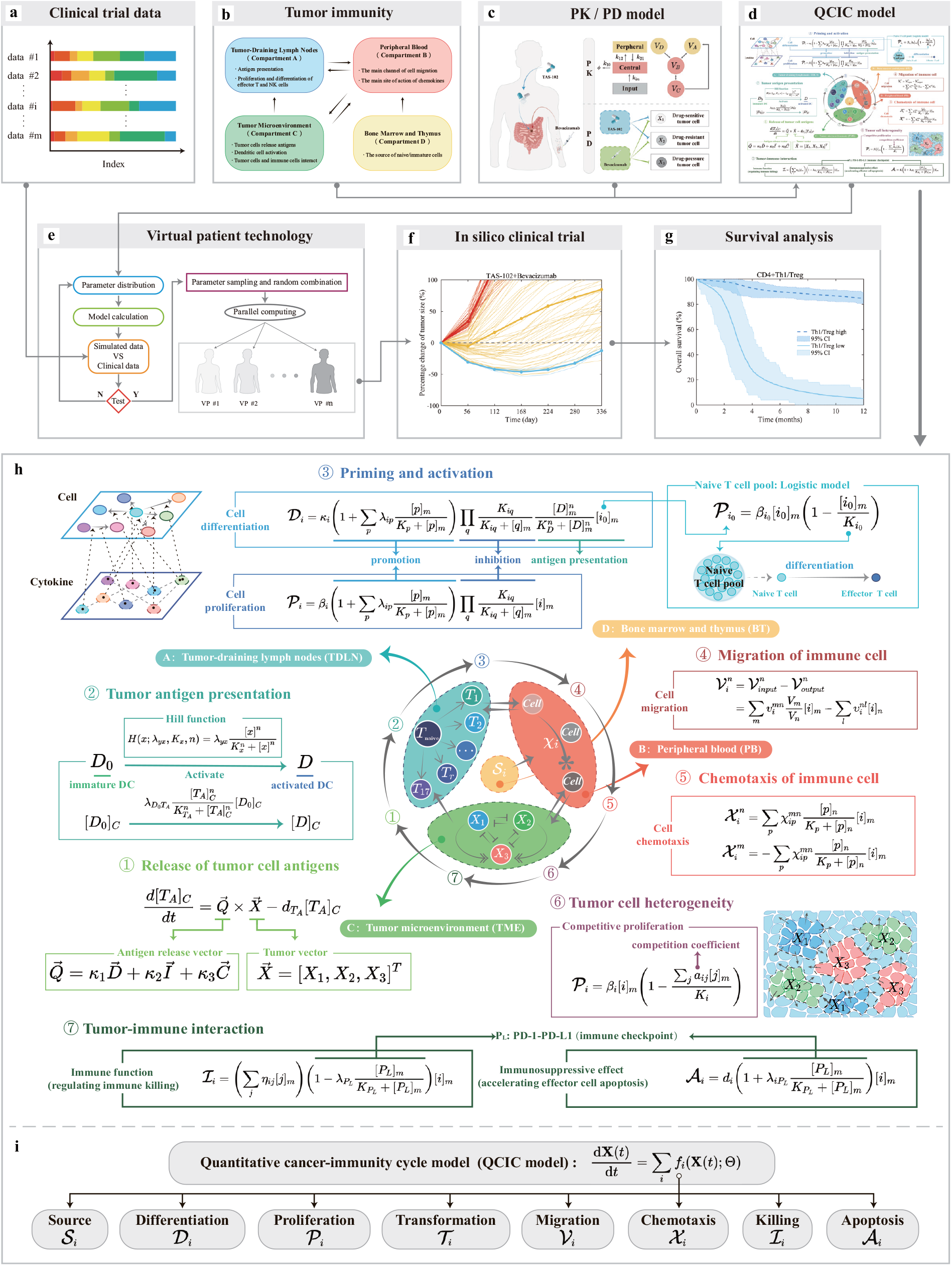
Illustration of the research framework for data-driven virtual patient generation and *in silico* clinical trials using the QCIC model. **a**. Clinical trial data-derived evaluation indexes of tumor treatment efficacy. **b**. Mechanisms of tumor immunity. **c**. Pharmacokinetic/pharmacodynamic (PK/PD) model. **d**. Integration of the PK/PD model with tumor tumor immunity mechanisms to form the QCIC model. **e**. Generation of virtual patients guided by clinical data using virtual patient technology. **f**. *In silico* clinical trials predicting tumor patient responses to various treatment options. **g**. Prediction of survival rates for different mCRC patients based on QCIC model-derived predictive biomarkers. **h**. The mathematical description of the seven main steps of the QCIC model. Step 1 involves tumor cell antigens release via mechanisms including normal apoptosis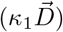), immune attack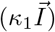, and chemotherapy killing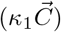. Step 2 depicts tumor antigen presentation facilitated by activated dendritic cells. Step 3 encompasses the priming and activation of T cells, with differentiation and proliferation described by the Michaelis-Menten and Hill functions, respectively. Steps 4 and 5 denote immune cell migration and chemotaxis. Step 6 represents tumor cell heterogeneity, including drug-sensitive (*X*_1_), drug-resistant (*X*_2_), and drug-pressure (*X*_3_) tumor cell types. Step 7 illustrates tumor-immune interactions, including immune cell targeting of tumor cells and tumor-induced immunosuppression via the PD-1-PD-L1 pathway. **i**. Symbolic representation of the QCIC model. For detailed descriptions of mechanisms, symbol meanings, and dynamic equations, please refer to the Methods section, Supplementary Text 1, and Supplementary Text 2.

The framework of our study is outlined as follows. Firstly, we selected 10 representative mCRC clinical trials with the same management and requirements between 2012 and 2023 (Fig. 1a) [55–64]. From these trials, we extracted evaluation indexes of clinical tumor efficacy, including complete response (CR), partial response (PR), stable disease (SD), progressive disease (PD), objective response rate (ORR), and disease control rate (DCR). These indexes were employed to assess the capability of the QCIC model to predict inter-individual treatment variances in mCRC patients.

Additionally, survival curves from various clinical trials were utilized to evaluate the accuracy of the QCIC model.

Secondly, to depict the entire process of the human immune response, the QCIC model was designed as a multi-compartmental ODE model according to the tumor immunity mechanisms (Fig. 1b). This model comprises four compartments: tumor-draining lymph node (TDLN), peripheral blood (PB), tumor microenvironment (TME), and the bone marrow and thymus (BT). Each compartment delineates distinct aspects of the immune response, with TDLN focusing on antigen presentation and cell differentiation, PB facilitating immune cell transport between tissues, TME representing the complex interplay between tumor and immune cells, and BT serving as the primary site for immune cell production.

Next, we integrated a pharmacokinetic (PK)/pharmacodynamic (PD) model into the QCIC model to illustrate the mechanism of drug action (Fig. 1c). In this context, the drug concentration in the central compartment corresponds to the concentration in tumor tissue. Thus, the QCIC model that couples the mechanism of drug action was developed (Fig. 1d).

Finally, model parameters were calibrated using virtual patient technology to generate a cohort reflecting clinical outcomes (Fig. 1e and 1f). During virtual patient generation, we primarily utilized six previously extracted clinical indexes (CR, PR, SD, PD, ORR, and DCR) to calibrate parameters describing tumor heterogeneity in the QCIC model. Consequently, our model captures inter-individual treatment differences and effectively correlates clinical treatment indexes with the QCIC model outputs.

Furthermore, tumor predictive biomarkers were extracted through binary classification of the virtual patient cohort, facilitating survival analysis on different subpopulations (Fig. 1g).

The biological mechanisms and mathematical formulas of the QCIC model are delineated below and depicted in Figure 1h, respectively.

1. Cancer cell antigen release: Within the tumor microenvironment (TME), tumor-associated antigens released by necrotic tumor cells are captured and processed by dendritic cells, initiating systemic immune responses [65].
2. Cancer cell antigen presentation: Dendritic cells, a key class of antigen-presenting cells (APCs), process antigens and present them as peptide major histocompatibility complex (p-MHC) molecular complexes [66–68].
3. T cell activation: Antigen-loads dendritic cells migrate to tumor-draining lymph node (TDLN) via the lymphatic circulation, where they present antigenic peptides to naïve T cells, promoting effector T cell production and proliferation [69, 70].
4. Trafficking of T cells to tumors: Effector T cells traverse the peripheral blood (PB) system to reach the vicinity of tumor tissue [71, 72].
5. T cell infiltration into tumors: Effector T cells from the peripheral blood infiltrate tumor, facilitated by vascular permeability and chemokine-mediated signaling [73].
6. T cells recognize cancer cells: Effector T cells within the TME specifically recognize and bind to cancer cells via the T cell receptor (TCR) [74].
7. Killing of cancer cells: Effector T cells execute tumor cell lysis through direct contact or release of cytotoxic substances such as granzyme, perforin, and interferon, thereby eliminating tumor lesions [75, 76].

The QCIC model considers the evolution dynamics of various types of tumor cells and immune cells. The formulation of the model accounts for the dynamic behavior of cell type *i* through several processes: source (*𝒮*_*i*_), differentiation (*𝒟*_*i*_), proliferation (*𝒫*_*i*_), transition (*𝒯*_*i*_), migration (*𝒱*_*i*_), chemotaxis (*𝒳*_*i*_), killing (*ℐ*_*i*_), and apoptosis (*𝒜*_*i*_). For instance, let [*i*]_*n*_(*t*) denotes the density of cell type *i* in compartment *n* at time *t*, the dynamical equation for [*i*]_*n*_(*t*) is given by (Fig. 1i)

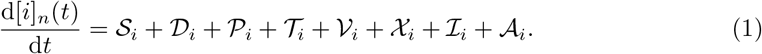

In the following, we present the mathematical description of different dynamic processes, respectively.

#### Cell source term: *𝒮*_*i*_

The source term is represented as:

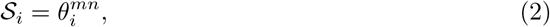

illustrates the release of cell type *i* from compartment *m* to compartment *n*. This term reflects the ongoing generation of naïve or immature cells originating from the bone marrow and thymus.

#### Cell differentiation term: *𝒟*_*i*_

Cell differentiation manifests in two forms. Firstly, naïve T cells within the tumor-draining lymph node (TDLN) differentiate into various effector T cell types under the influence of activated DCs and cytokines. Mathematically, the differentiation process of naïve T cells is characterized by a combination of the Michaelis-Menten function and Hill function [77–79]

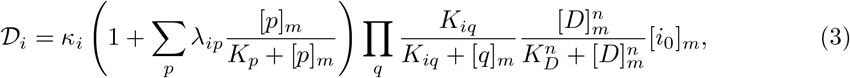

where *κ*_*i*_ denotes the differentiation rate of cell type *i*; λ_*ip*_ signifies the enhancement coefficient of cytokine *p* on the differentiation process of cell type *i*; [*p*]_*m*_ and [*q*]_*m*_ represent the concentrations of cytokines *p* and *q* in compartment *m*, respectively; *K*_*p*_ indicates the half-saturation constant of cytokines *p*; *K*_*iq*_ denotes the inhibition of cell type *i* function by cytokine *q*; *K*_*D*_ represents the half-saturation constant of activated DCs; *n* represents the Hill’s coefficient during the antigen presentation; [*D*]_*m*_ represents the density of activated DCs in compartment *m*.

Secondly, neutrophils and macrophages within the tumor microenvironment (TME) undergo further polarization into two phenotypes: pro-tumor and anti-tumor.

Polarization, akin to differentiation, occurs at a low rate and is confined to specific cells. The mathematical representation of polarization is expressed as:

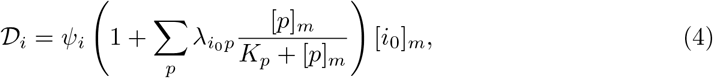

where *ψ*_*i*_ signifies the polarization rate from cell type *i*_*0*_ to cell type *i*, with *i*_*0*_ representing the precursor of cell type *i*; 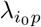 denotes the enhancement coefficient of cytokine *p* on the polarization process of cell typpe *i*_*0*_.

#### Cell proliferation term: 𝒫_*i*_

The cell proliferation term is delineated in three distinct forms. Firstly, the proliferation of naïve T cells in TDLN adheres to the logistic equation as [78]:

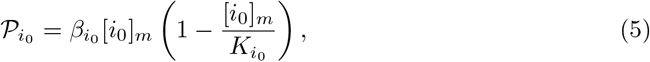

where 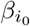 signifies the proliferation rate of naïve T cells; 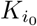 indicates the carrying capacity of naïve T cells.

Secondly, lymphocytes, which encompass various T cell subpopulations and NK cells in the TDLN, are not terminally differentiated cells but rather proliferative. Their proliferation terms under cytokine influence are described as follows:

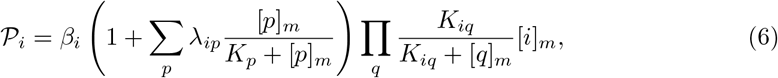

where *β*_*i*_ denotes the proliferation rate of cell *i*; λ_*ip*_ represents the enhancement coefficient of cytokine *p* on the proliferation of cell *i*; *K*_*p*_ represents the half-saturation constant of cytokines *p*; *K*_*iq*_ denotes the inhibition of cell type *i* function by cytokine *q*.

Finally, for the three tumor cell types–drug-sensitive tumor cells (DSTC), drug-resistant tumor cells (DRTC), and drug-pressure tumor cells (DPTC)–their proliferation terms adhere to competing logistic equations [80–82]:

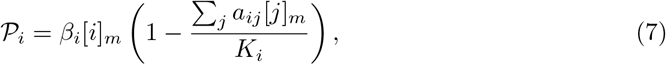

where *K*_*i*_ represents the carrying capacity of tumor cell *i*; *a*_*ij*_ signifies the intraspecific competition coefficient (*i* = *j*) or the interspecific competition coefficient (*i≠ j*).

#### Cell type transition term: *𝒯*_*i*_

Cell type transition may occur under the influence of cytokines or due to drug pressures, reflecting cellular plasticity. Cell type transition can be mathematically represented as [32]:

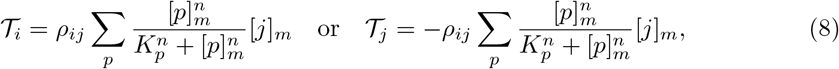

where *ρ*_*ij*_ represents the transition rate from cell type *j* to cell type *i*; [*p*]_*m*_ represents the concentrations of cytokines or drugs *p* in compartment *m*; *K*_*p*_ represents the half-saturation constant of cytokines or drugs *p*; *n* represents the Hill’s coefficient during cell phenotype transition. Specifically, *n* = 1 is selected for cytokine *p*, and *n* > 1 for drugs *p*.

#### Cell migration term: *𝒱*_*i*_

Cell migration between different compartments is represented as [78, 83]:

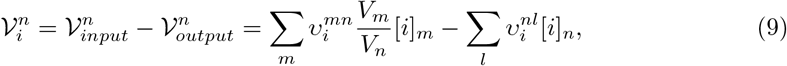

where 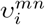 and 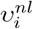 represent the migration rate of cell type *i* from compartment *m* to compartment *n*, and the migration rate of cell type *i* from compartment *n* to compartment *l*; *V*_*m*_ and *V*_*n*_ represent the volume of compartments *m* and *n*, respectively.

#### Cell chemotaxis term: *𝒳*_*i*_

The cell chemotaxis term is represented as [43, 44]

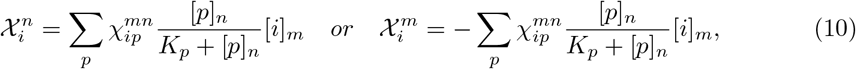

where 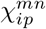 represents the average chemotactic rate of cell type *i* from compartment *m* to compartment *n* under the action of chemokine *p*; *K*_*p*_ represents the half-saturation constant of chemokine *p*.

#### Cell killing term: *ℐ*_*i*_

The cell killing term is represented as [32, 34, 36, 84]:

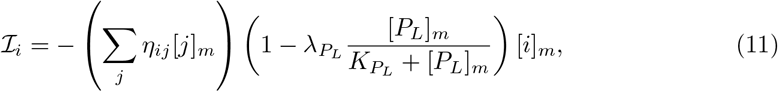

where *η*_*ij*_ represents the immune killing of tumor cell type *i* by immune cell type 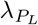 represents the killing regulation coefficient of tumor by PD-1-PD-L1; 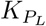 represents the half-saturation constant of PD-1-PD-L1; [*P*_*L*_]_*m*_ represents the concentrations of PD-1-PD-L1 in compartment *m*.

#### Cell apoptosis term: *𝒜*_*i*_

According to the mechanism of cell apoptosis, the cell apoptosis term is primarily represented in three forms. Firstly, activation-induced cell death (AICD) refers to the mechanism by which activated lymphocytes can undergo self-limited cell death through the interaction between Fas and FasL [85, 86]:

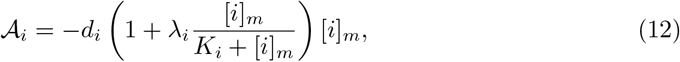

where *d*_*i*_ represents the apoptosis rate of cell type *i*; λ_*i*_ represents the apoptosis regulatory coefficient caused by the self-limiting of cell type *i. K*_*i*_ represents the half-saturation constant of cell type *i*.

Secondly, tumor cells can induce programmed death of T cells and NK cells by PD-1-PD-L1 [87–89]:

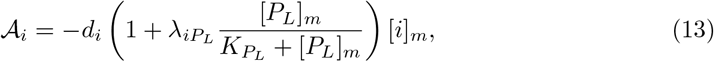

where *d*_*i*_ represents the apoptosis rate of cell type 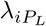 represents the apoptosis regulation coefficient of cell type *i* by PD-1-PD-L1.

Finally, apoptosis induced by chemotherapy can be expressed as [90–92]:

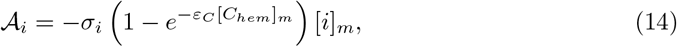

where *σ*_*i*_ represents the killing rate of cell type *i* by chemotherapy; ε_*C*_ represents the effective coefficient of chemotherapy; [*C*_*hem*_]_*m*_ represents the concentration of the chemotherapy drug in compartment *m*.

Additionally, the dynamic changes at the molecular level can be regarded as fast-scale behaviors compared to the dynamic evolution at the cellular level. Therefore, the dynamic behavior of cytokine or chemokine *p* in compartment *n* is described as

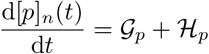

where 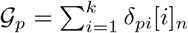 and *ℋ*_*p*_ = *−d*_*p*_[*p*]_*n*_ denote cytokine or chemokine production and degradation, respectively; *k* represents the number of cell types that can secrete cytokine or chemokine *p*; δ_*pi*_ represents the cytokine or chemokine *p* secretion rate by cell type *i*, and *d*_*p*_ represents the degradation rate of cytokine or chemokine *p*.

Based on the construction principles of the QCIC model, we constructed an ODE model with 95 variables to describe the dynamic process of the tumor-immune response. Specific details regarding the dynamics of tumor cells, immune cells, cytokines, and chemokines are described in the Methods section and Supplementary Text 1.

### Prediction of short-term treatment efficacy in advanced mCRC patients

In clinical trials, short-term treatment efficacy of the drugs in advanced mCRC patients is often assessed using six key indices: complete response (CR), partial response (PR), stable disease (SD), progressive disease (PD), objective response rate (ORR), disease control rate (DCR), with ORR defined as CR + PR and DCR as CR + PR + SD. These evaluation indexes are determined based on changes in tumor status before and after treatment [93]. In this study, we analyzed data from 10 clinical trials, integrating information from 2,566 patients with advanced mCRC (Supplementary Figure 1) [55–64]. Given the poor prognosis of advanced mCRC patients ineligible for surgery, CR, PR, and ORR are rarely observed, making SD, PD, and DCR the most pertinent short-term efficacy indices.

To validate the predictive capability of the QCIC model for short-term outcomes in advanced mCRC patients, we conducted *in silico* clinical trials by generating 100 virtual patients using virtual patient technology and compared the model predictions with clinical data. We introduced the treatment response index (TRI) to quantify the results of virtual clinical trials:

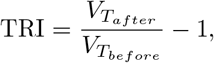

where 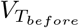 represents the pre-treatment tumor volume (baseline), and 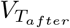 represents post-treatment tumor volume. TRI primarily reflects changes in tumor volume relative to baseline after treatment. According to Response Evaluation Criteria in Solid Tumours (RECIST) version 1.1 [93], patients were classified as PD if TRI ≥ 0.2, otherwise as DCR, and as SD if the TRI ≥ −0.3 and < 0.2. This allows translation of *in silico* trial results into clinical metrics such as SD, PD, and DCR.

Model simulations indicated that patients with advanced mCRC treated with placebo exhibited SD, PD, and DCR rates of 15%, 85%, and 15%, respectively (Fig. 2a), closely matching clinical trial data: 15.3%, 84.5%, and 15.5%. Similarly, patients receiving TAS-102 chemotherapy showed SD, PD, and DCR rates of 44% vs. 44.4%, 55% vs. 54.3%, and 45% vs. 45.7% in model simulations vs. clinical trials, respectively (Fig. 2b). Furthermore, combination therapy with TAS-102 plus bevacizumab resulted in SD, PD, and DCR rates of 61% vs. 58.9%, 36% vs. 37.2%, and 64% vs. 62.8% in model simulations vs. clinical trials, respectively (Fig. 2c). These results demonstrate the ability of the QCIC model to accurately predict short-term treatment efficacy in advanced mCRC patients. A comparison of model simulations with different clinical trials is provided in Supplementary Figure 2.

**Fig 2.**
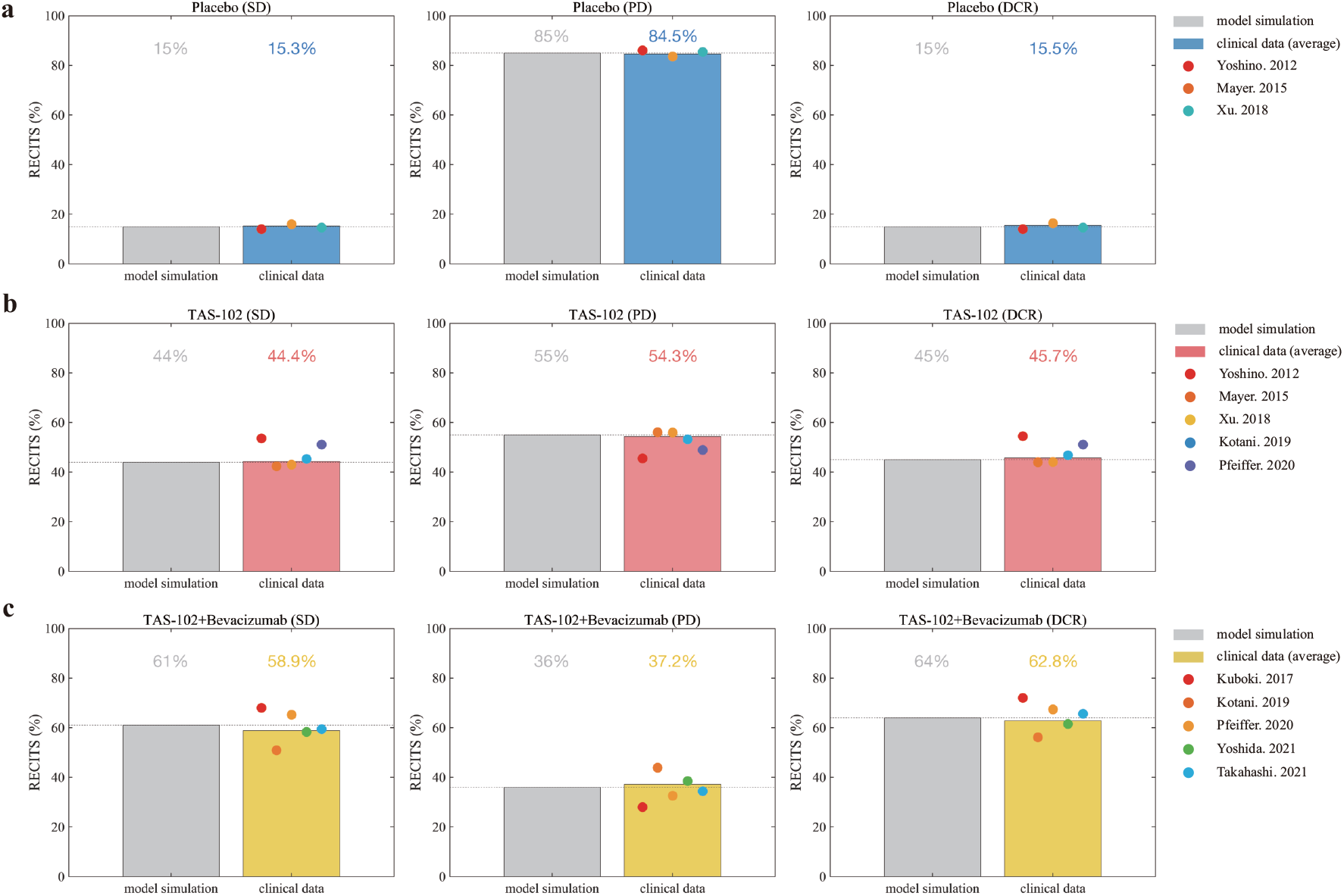
Validation of the QCIC model at the population level by comparing short-term treatment efficacy evaluation indexes (SD, PD, and DCR) from model simulations and clinical data. **a**. Placebo group. **b**. TAS-102 group. **c**. TAS-102+bevacizumab group. Bar graphs display the results of the model simulations and the averaged clinical data. Different colored dots represent different clinical data. The dashed lines show the results of the model simulations used to observe the variations across different clinical trials.

To further elucidate the QCIC model’s role in predicting cancer progression, we depicted inter-individual treatment differences among patients captured by virtual clinical trials and treatment outcomes at the initial follow-up (eighth week) using spider plots (left) and waterfall plots (right) in Fig.3, respectively. DCR increased from 15% to 45% in patients receiving TAS-102 chemotherapy (Fig. 3a and b), indicating that 30% of advanced mCRC patients may benefit significantly from TAS-102 chemotherapy.

**Fig 3.**
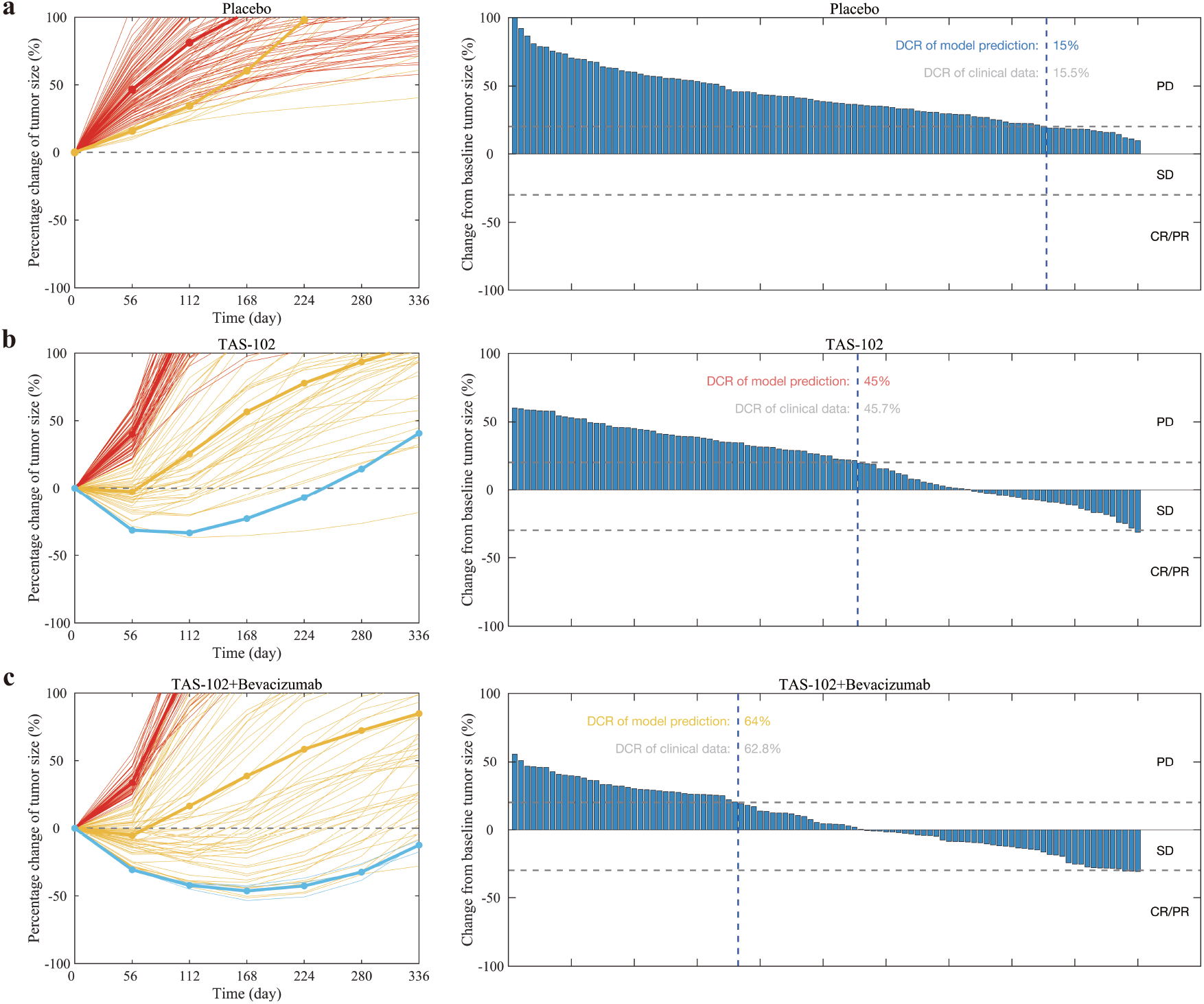
Virtual patients were generated using the QCIC model to predict dynamic tumor evolution (left) and short-term efficacy (right). Treatment response is evaluated based on RECIST V.1.1 in the placebo group (**a**), TAS-102 group (**b**), and TAS-102+bevacizumab group (**c**). Tumor volume change was recorded every two treatment cycles (every eight weeks), consistent with medical imaging used to diagnose disease progression during clinical treatment. In spider plots on the left, each thin line represents a virtual patient, and the thick line represents their average performance. Red denotes PD patients, yellow denotes SD patients, and blue denotes CR/PR patients. In the waterfall plot on the right, each bar represents tumor volume change relative to baseline for a virtual patient during the first imaging examination (eighth week). The left side of the blue dotted line denotes the PD patient population, and the right side denotes the DCR patient population.

These findings suggest the ability of TAS-102 chemotherapy to notably improve tumor burden in advanced mCRC patients. Furthermore, combination therapy with TAS-102 plus bevacizumab increased DCR from 15% to 64% and reduced disease progression from 85% to 36% (Fig. 3a and c), indicating nearly 50% of patients may significantly benefit from combination therapy. This suggests combination therapy with bevacizumab improved DCR by 20% in advanced mCRC patients compared to TAS-102 monotherapy. Thus, model simulations unveil additional clinical benefits associated with combination therapy.

### Prediction of overall survival in advanced mCRC patients

To delve into the capacity of the QCIC model in predicting the long-term efficacy of drug therapy in advanced mCRC patients, we introduce a death probability function (DPF) based on tumor burden. The DPF is expressed as

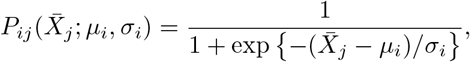

where *i* represents the *i*-th type of treatment option, *µ*_*i*_ and *σ*_*i*_ represent the shape parameters of the *i*-th type of treatment option, and 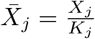 represents the ratio of the dominant tumor cell type *X*_*j*_ to the carrying capacity *K*_*j*_. *P*_*ij*_ describes the death probability caused by the dominant tumor cell type *X*_*j*_ under the *i*-th treatment option. Our model accounts for three tumor cell types: drug-sensitive tumor cells (DSTC, *X*_1_), drug-resistant tumor cells (DRTC, *X*_2_), and drug-pressure tumor cells (DPTC, *X*_3_). To ensure robustness, we generated 4000 virtual patients and randomly selected 500 patients for *in silico* clinical trials.

Results showed the median overall survival (M-OS) for advanced mCRC patients treated with placebo was 6.1 months, closely aligned with clinical outcomes (6.01 months) (Fig. 4a). Similarly, for TAS-102 chemotherapy, M-OS was 7.6 months in simulations vs. 7.52 months clinically (Fig. 4a). Additionally, TAS-102 plus bevacizumab treatment resulted in an M-OS of 10.1 months in simulations, closely matching clinical data (10.01 months) (Fig. 4a). These findings show that the QCIC model is capable of predicting the long-term outcomes after drug therapy.

**Fig 4.**
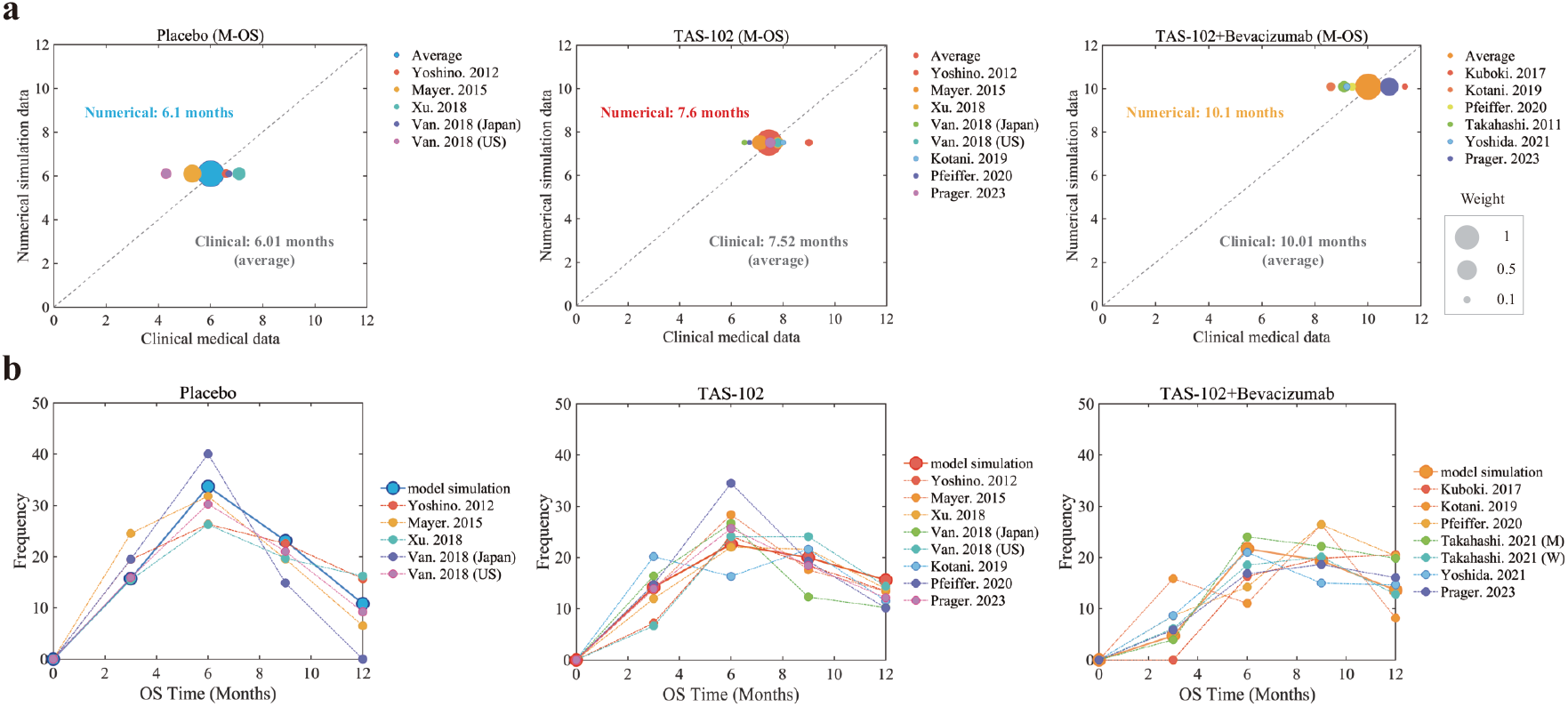
Comparison of long-term efficacy evaluation indices in model simulations and clinical data. **a**. Median overall survival (M-OS) comparison between model simulations and clinical data. Different color dots represent results from various clinical trials, with dot size weighted by the number of patients. **b**. Validation of QCIC model prediction at the population scale by comparing the predicted death frequency with clinical death data at the 3rd, 6th, 9th, and 12th months post-treatment. The thick solid line depicts model-calculated results, while the dashed line represents clinical trial outcomes.

To further explore the predictive capability of the QCIC model for long-term outcomes, we examined how treatment affected death frequency by comparing model predictions with clinical data over 3, 6, 9, and 12 months post-treatment. Our analysis revealed a significant decrease in patient mortality with TAS-102 chemotherapy compared to the placebo group, particularly evident at the 6-month mark, decreasing from 33.7% to 22.6% (Fig. 4b). Furthermore, combination therapy substantially reduced patient deaths within the initial 3 months, dropping from 14.2% to 4.7% (Fig. 4b). These findings underscore the efficacy of TAS-102 chemotherapy in increasing the six-month survival rate among advanced mCRC patients compared to those receiving placebo treatment. Additionally, combination therapy of TAS-102 plus bevacizumab further improves the survival rate in the early treatment phase (within 3 months). Model predictions depicted evolution dynamics under treatment consistent with the corresponding clinical data, indicating that model simulation can provide a confident prediction of long-term outcomes.

To track disease progression in advanced mCRC patients, we depicted overall survival under different treatment options using Kaplan-Meier (K-M) survival curves, integrating both clinical data and simulation results (Fig. 5a-c). Additionally, Figure 5d presents the overall survival over time of advanced mCRC patients under varied treatments. Notably, TAS-102 chemotherapy extended median overall survival (M-OS) from 6.1 to 7.6 months, prolonging it by 1.5 months. Furthermore, TAS-102 plus bevacizumab boosted M-OS by nearly 2.5 months, achieving 10.1 months in advanced mCRC patients. From a one-year overall survival standpoint, the combination therapy of TAS-102 plus bevacizumab enhanced the patient’s survival by almost 24%, increasing from 16% to nearly 40% (Fig. 5d). TAS-102 monotherapy improved one-year overall survival by almost 11%, and adding bevacizumab to combination therapy can elevate the one-year overall survival by 13% (Fig. 5d).

**Fig 5.**
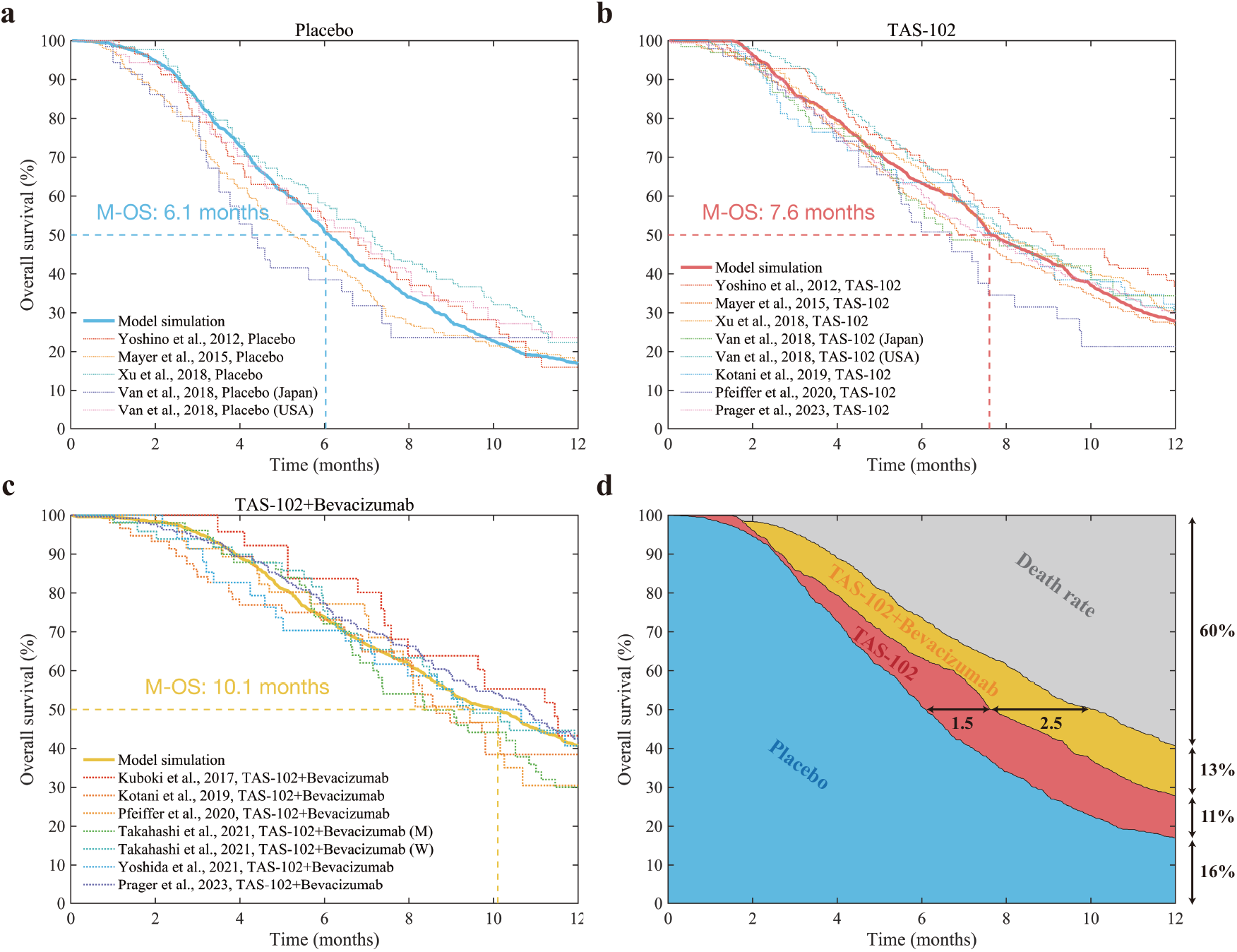
Overall survival of advanced mCRC patients on the population scale. **a**. Placebo group. **b**. TAS-102 group. **c**. TAS-102+bevacizumab group. **d**. Comparison of overall survival in advanced mCRC patients across various treatment options. In the figure, the solid think lines in blue, red, and orange represent the survival curves predicted by the model for the placebo group, TAS-102 chemotherapy group, and TAS-102+bevacizumab group, respectively. The other dashed lines represent survival curves from different clinical trials.

### Predictive biomarkers for mCRC based on the QCIC model

To further explore the role of tumor-infiltrating lymphocytes (TILs) in the immune response, we examined four indices: the density of tumor-infiltrating CD4+Th1 cells, the density of tumor-infiltrating CD8+CTL cells, the tumor-infiltrating CD4+Th1/Treg ratio, and the tumor-infiltrating CD8+CTL/Treg ratio, as potential biomarkers for predicting disease progression in advanced mCRC patients. Using data from 500 virtual patients generated from *in silico* clinical trials, we classified them as responders (R) or non-responders (NR), defining responders as CR+PR+SD patients and non-responders as PD patients due to the low occurrence of CR and PR in advanced mCRC. We then compared the distribution of these predictive biomarkers between responders and non-responders.

The analysis revealed that the density of tumor-infiltrating CD8+ CTL cells exhibited greater variability between responders (R) and non-responders (NR) compared to the density of tumor-infiltrating CD4+Th1 cells (Fig. 6). This suggests that the density of tumor-infiltrating CD8+CTL cells may be a crucial predictive index. Additionally, the tumor-infiltrating CD8+CTL/Treg displayed less variability between responders (R) and non-responders (NR) compared to the tumor-infiltrating CD4+Th1/Treg ratio (Fig. 6), indicating that the former may not be a strong predictive index. Despite differences in performance among the various indices, the results consistently showed a higher abundance of tumor-infiltrating immune cells in responders (NR).

**Fig 6.**
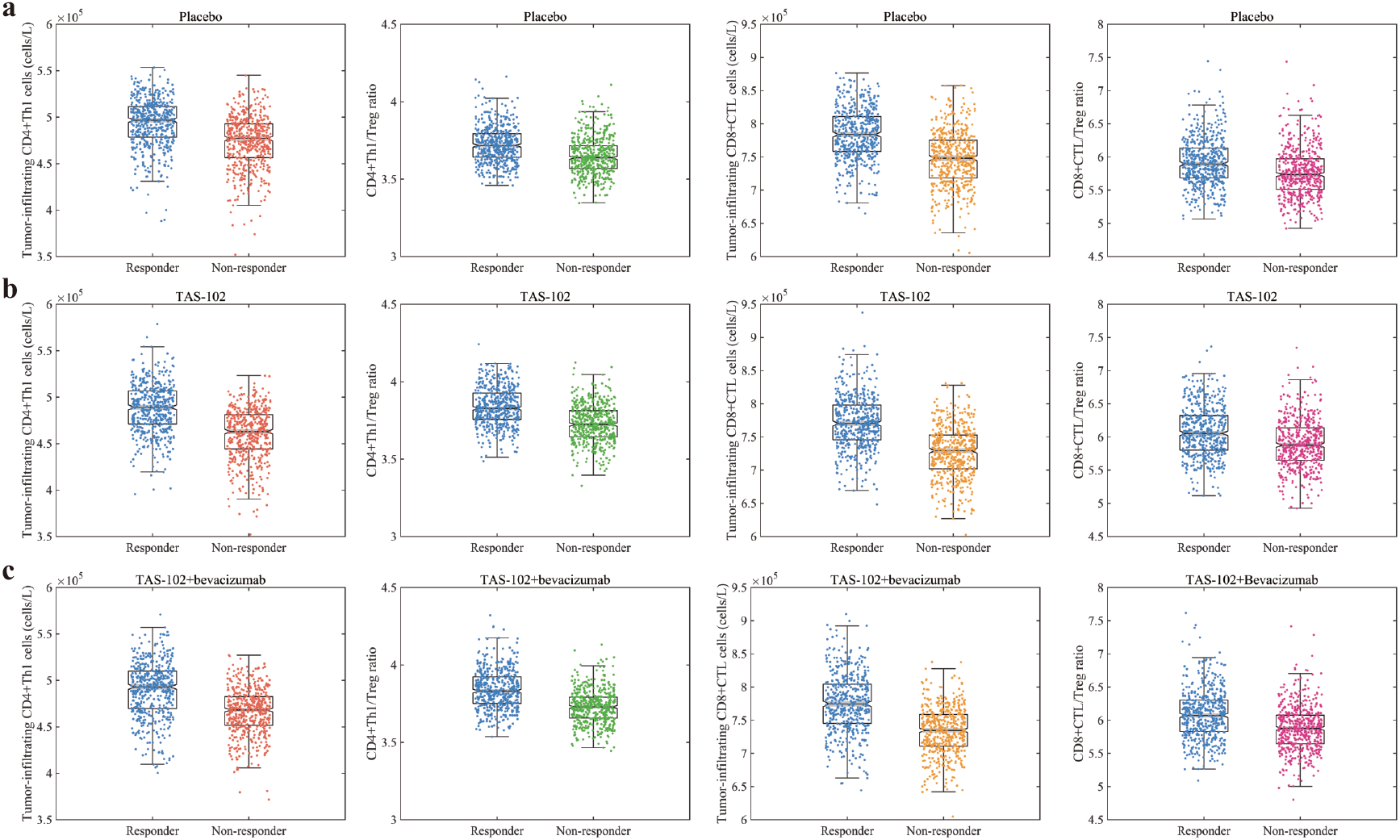
Distribution of predictive biomarkers calculated by QCIC model in responders and non-responders. **a**. Placebo group. **b**. TAS-102 group. **c**. TAS-102+bevacizumab group. The data samples are visually represented by box plots, where the line inside each box indicates the median of the sample, and the upper and lower edges of each box represent the upper and lower quartiles, respectively. The p-value of both sets of data in each subplot is less than 0.001 (t-test).

To further evaluate the performance of these predictive biomarkers, we constructed receiver operating characteristic (ROC) curves using machine learning binary classification models. We used numerical simulation data from the initial clinical diagnosis (the eighth week) as the test set to assess the predictive ability of these biomarkers for short-term treatment outcomes. The median value of each biomarker across 500 virtual patients served as the threshold for ROC analysis. As depicted in Figure 7, the density of tumor-infiltrating CD8+CTL cells exhibited the highest area under the curve (AUC) in both TAS-102 monotherapy and combination therapy with bevacizumab, reaching 0.77 (Fig. 7). Hence, the density of tumor-infiltrating CD8+CTL cells emerges as a critical predictive biomarker for short-term treatment efficacy in advanced mCRC patients. Conversely, the tumor-infiltrating CD8+CTL/Treg ratio demonstrated the poorest predictive performance among the four biomarkers (Fig. 7).

**Fig 7.**
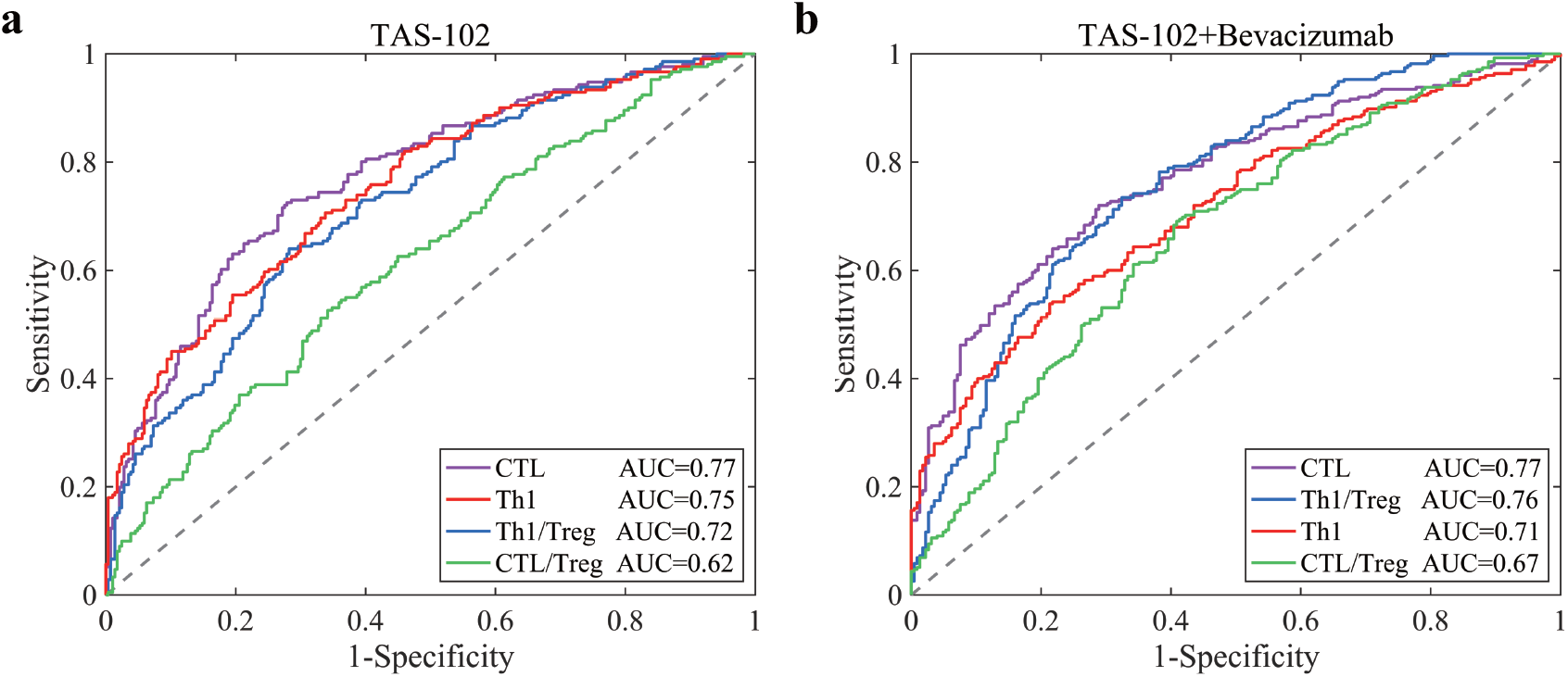
ROC analysis of predictive biomarkers in TAS-102 monotherapy and combination therapy with bevacizumab. **a**. TAS-102 group. **b**. TAS-102+bevacizumab group. The response status (R vs NR) for each virtual patient was predicted by comparing the pretreatment levels of predictive biomarkers to the cut-off value.

### Survival prognosis based on predictive biomarkers of mCRC

To enhance the survival outcomes of advanced mCRC patients, we delved into predictive biomarkers for prognostic analysis during clinical treatment. We divided the 500 virtual patients into high- and low-expression groups based on predictive biomarker levels. Subsequently, we assessed patient survival using DPF, and Figure 8 illustrates the K-M survival curves.

**Fig 8.**
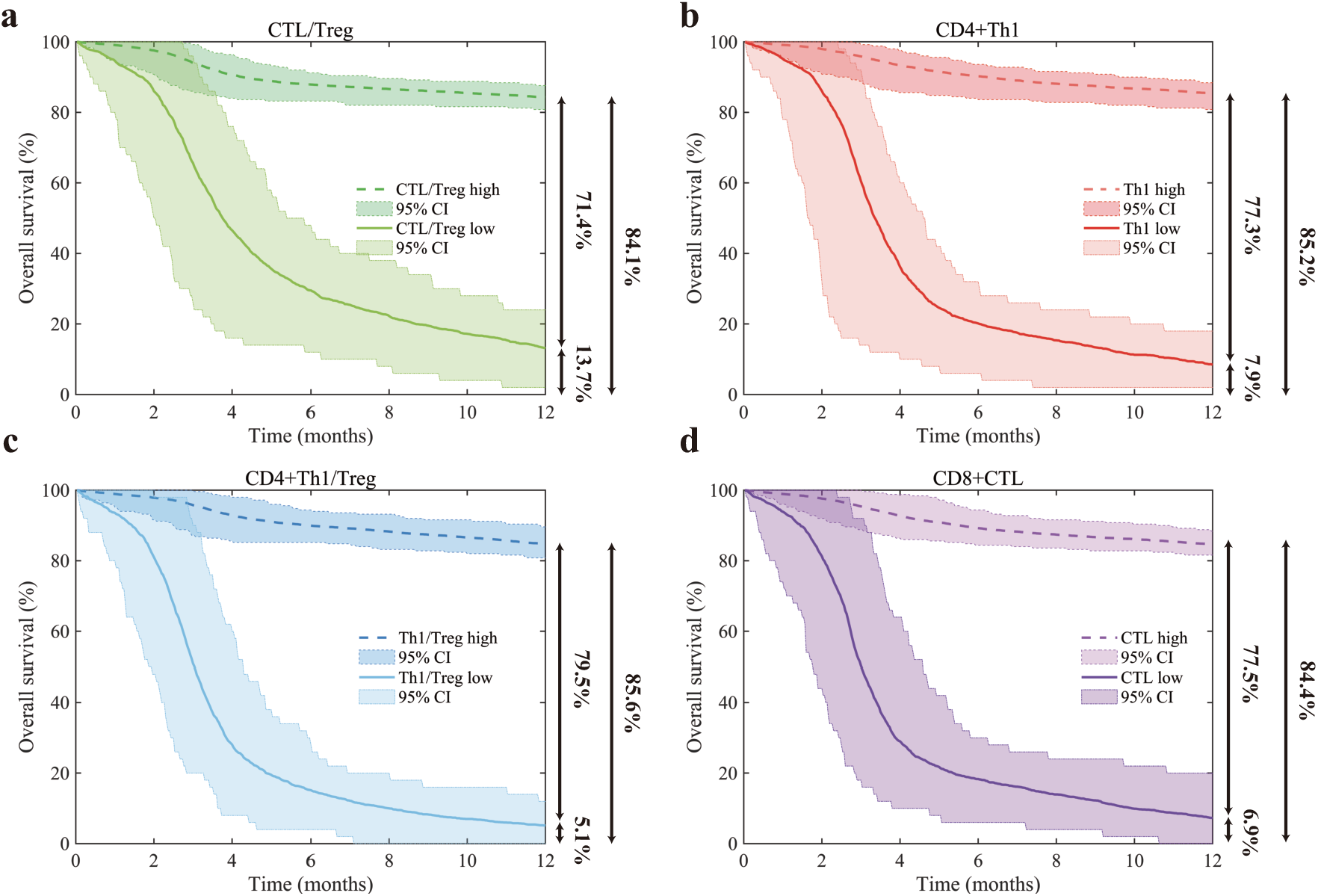
Prognostic analysis of predictive biomarkers in combination therapy with TAS-102 plus bevacizumab. Panels **a** through **d** sequentially illustrate the overall survival analysis in patients with advanced mCRC based on four predictive biomarkers: the tumor-infiltrating CD8+CTL/Treg ratio, the density of tumor-infiltrating CD4+Th1 cells, the tumor-infiltrating CD4+Th1/Treg ratio, and the density of tumor-infiltrating CD8+CTL cells. In each panel, the dashed line represents the high expression of the predictive indicator, the solid line represents the low expression, and the shaded area indicates the 95% confidence interval.

Our findings revealed that all four predictive biomarkers significantly influenced patient survival prognosis. However, distinct variations in survival prognosis were observed among different biomarkers. Notably, a lower tumor-infiltrating CD8+CTL/Treg ratio demonstrated superior survival prognosis in the trial group with low expression of all predictive biomarkers, with a one-year overall survival rate of 13.7% (Fig. 8a). Moreover, the 95% confidence intervals for the tumor-infiltrating CD8+CTL/Treg ratio and the density of tumor-infiltrating CD4+Th1 cells exhibited broader ranges, indicating greater fluctuations in both indices (Fig. 8a and b).

The tumor-infiltrating CD4+Th1/Treg ratio emerged as the most significant indicator of survival difference between high and low expression groups for the same predictive biomarker (Fig. 8c). It showcased a substantial 79.5% difference in one-year overall survival (Fig. 8c). Similarly, the density of tumor-infiltrating CD8+CTL cells displayed a strong predictive capability, with a 77.5% difference in one-year overall survival, albeit slightly less significant than the tumor-infiltrating CD4+Th1/Treg ratio (Fig. 8d). In summary, the tumor-infiltrating CD4+Th1/Treg ratio emerged as the most remarkable clinical index for predicting survival prognosis in advanced mCRC patients, as indicated by the QCIC model we developed.

## Discussion

Mathematical modeling of tumor treatments has become a critical area in mathematical oncology and computational systems biology, with numerous studies highlighting its significance [13–28]. Various mathematical methods, including response-diffusion equations, have been applied to predict the dynamic evolution of tumors during combination therapy by quantifying inter-regulatory relationships between tumor cells, immune cells, and cytokines in the tumor microenvironment [29–37]. However, the systematic development of mathematical models accurately describing treatment differences between patients, particularly predictive models for disease heterogeneity during tumor treatment, is still in its early stages. Recent advances in quantitative systems pharmacology (QSP) models based on pharmacokinetic mechanisms have addressed inter-individual treatment differences and tumor recurrence phenomena in cancer patients [38–50]. Nevertheless, analyzing the survival prognosis of tumor patients using predictive biomarkers extracted from mathematical models remains challenging and represents an important area for future research.

Patients with advanced mCRC exhibit significant inter-individual heterogeneity in clinical treatment responses. A quantitative understanding of the interaction between tumor evolution dynamics and the host immune response can adequately capture inter-individual treatment differences. In this study, we developed the quantitative cancer-immunity cycle (QCIC) model, a multi-compartment, multi-scale, multi-dimensional mathematical model, to quantitatively describe the main biological processes of the tumor immune response (Fig.1). The QCIC model integrates cellular-level biological mechanisms such as cell regeneration, differentiation, proliferation, apoptosis, migration, and chemotaxis, as well as molecular-level processes including antigen release, presentation, cytokine/chemokine secretion, degradation, and drug metabolism. By incorporating compartments for various biological sites, including the tumor microenvironment, peripheral blood, tumor-draining lymph nodes, bone marrow, and thymus, the model provides explicit spatial and temporal dimensions for immune response behaviors, capturing the heterogeneity observed in patients’ clinical responses. The QCIC model offers a systematic approach to inferring tumor evolution and inter-individual treatment differences.

Our study demonstrated that the QCIC model accurately derived short-term clinical efficacy evaluation indices consistent with the clinical trials (Fig. 2) and predicted the dynamic evolution of tumor burden in advanced mCRC patients (Fig. 3). Moreover, the model effectively assessed the long-term clinical outcomes, showing significant improvements in six-month survival rates with TAS-102 chemotherapy and reduced risk of early death with combination therapy of TAS-102 plus bevacizumab (Fig. 4).

Importantly, the model’s simulation of median overall survival aligned well with clinical data, indicating its capability to capture both short- and long-term clinical outcomes in mCRC patients (Fig. 5). Furthermore, we analyzed predictive biomarkers based on disease progression obtained from the QCIC model (Fig. 6), highlighting the tumor-infiltrating CD8+CTL cell density as a critical predictive biomarker for short-term treatment outcomes (Fig. 7). Additionally, prognostic analysis of patient survival revealed the tumor-infiltrating CD4+Th1/Treg ratio as a significant predictor of survival prognosis (Fig. 8). These findings demonstrate that the QCIC model bridges mathematical models with clinical prognosis, extending the research of existing models into clinical applications.

It is important to note that valid patients for the QCIC model are generated using clinical data on solid tumor evolution assessed through various imaging techniques.

These data calibrate the key parameters reflecting tumor heterogeneity in the model, providing a valuable reference for individualized treatment. The model’s ability to predict tumor dynamics after treatment based on pre-treatment medical data can guide timely adjustments to treatment strategies during clinical care. Although this study focused on the efficacy of TAS-102+bevacizumab therapy, the QCIC model can explore the benefits of other drug combinations with sufficient clinical data, offering patients optimal treatment options.

Despite the advancements, the QCIC model has limitations, such as incomplete capture of inter-cellular interactions and the inability to simulate inter-cell communications based on cell positional information. Future iterations may incorporate insights from recent studies on tumor immune dynamics, spatial heterogeneity of the tumor microenvironment, and drug action mechanisms. Additionally, stochastic simulation algorithms, machine learning methods, and agent-based models can enhance the QCIC model’s numerical simulation capabilities.

In conclusion, integrating mathematical models, machine learning, and virtual experiments provides a valuable approach to addressing inter-individual heterogeneity in clinical treatment responses. Predictive biomarkers extracted from mathematical models offer insights into the survival prognosis of cancer patients. Multidisciplinary studies, like the one presented here, contribute to understanding tumor evolution and therapeutic strategies, advancing the field of solid tumor treatment. The discoveries and methodology of this study lay a solid foundation for future research aimed at identifying optimal treatment combinations and improving solid tumor treatment options.

## Methods

### Clinical trials and data analysis

In this study, we selected 10 pivotal clinical trials conducted between 2012 and 2023, which employed consistent treatment regimens [55–64]. Detailed information regarding these trials is provided in Supplementary Figure 1, Supplementary Table 1, and Supplementary Table 2. Patients in the monotherapy received TAS-102 (35 mg/m^2^) orally twice daily on days 1-5 and days 8-12, with a 28-day cycle. Those assigned to combination therapy also received intravenous bevacizumab (5 mg/kg) on days 1 and 15. We meticulously integrated data from 2566 clinical patients, comprising 507 patients in the TAS-102 plus bevacizumab group, 1486 in the TAS-102 group, and 573 in the control group. To ensure the accuracy and utility of the clinical data, we excluded any unevaluated sections from the original dataset.

**Table 1.** Data analysis based on 10 clinical trials.

| Group | Number | CR | PR | SD | PD | ORR | DCR | M-OS |
| --- | --- | --- | --- | --- | --- | --- | --- | --- |
| TAS-102+Bevacizumab | 507 | 0% | 3.9% | 58.9% | 37.2% | 3.9% | 62.8% | 10.01 months |
| TAS-102 | 1486 | 0% | 1.3% | 44.4% | 54.3% | 1.3% | 45.7% | 7.52 months |
| Placebo | 573 | 0.2% | 0% | 15.3% | 84.5% | 0.2% | 15.5% | 6.01 months |

Explanation: complete response (CR), partial response (PR), stable disease (SD), progressive disease (PD), objective response rate (ORR), disease control rate (DCR), median overall survival (M-OS).

**Table 2.**
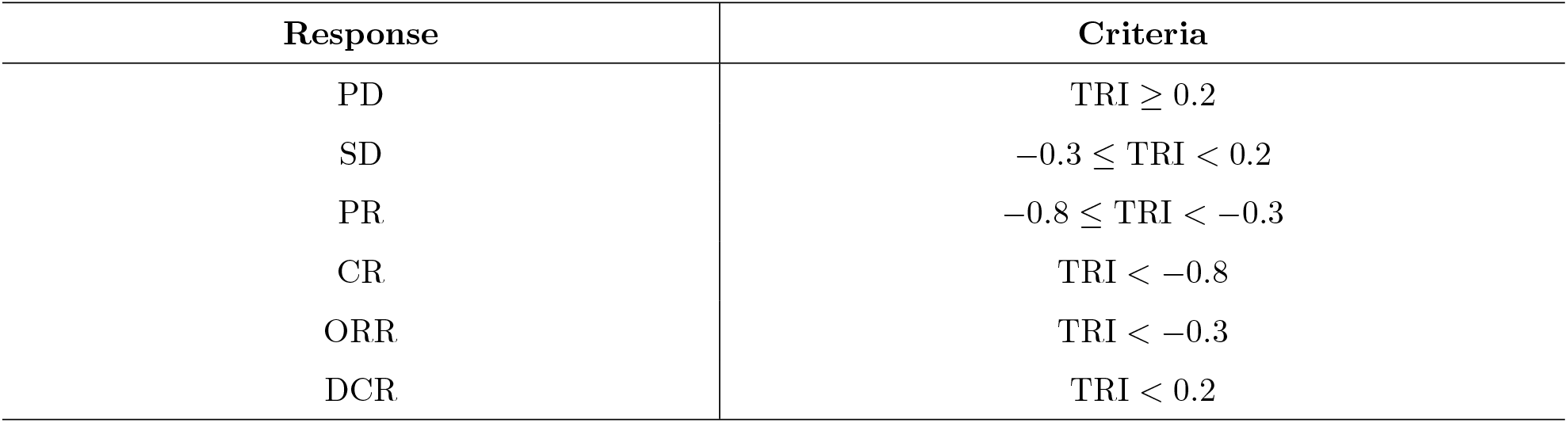
The dynamic evolution of solid tumors based on TRI was determined in virtual patients.

To synthesize the results from various clinical trials, we introduce the equation:

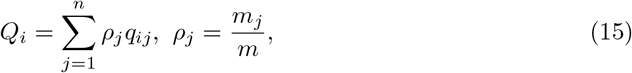

where *Q*_*i*_ represents different clinical indexes (*e*.*g*., CR, PR, SD, PD, ORR, DCR, and M-OS), *j* denotes different clinical trials, *ρ*_*j*_ signifies the weight assigned to the *j*-th clinical trial, *m* indicates the total number of patients, *m*_*j*_ represents the number of patients in the *j*-th clinical trial, and *q*_*ij*_ denotes the data of the *i*-th clinical index in the *j*-th clinical trial. The aggregated data is summarized in Table 1.

### The biological mechanisms of tumor immunity

The human body’s tumor immune response is divided into four compartments: tumor-draining lymph nodes (compartment A), peripheral blood (compartment B), tumor microenvironment (compartment C), and bone marrow and thymus (compartment D) (Fig.9a).

Tumor-draining lymph nodes (compartment A) are pivotal secondary lymphoid organs in tumor immunity, facilitating lymphocyte proliferation, differentiation, and activation of specific immunity [69, 70]. Within these lymph nodes, antigen-presenting cells (APCs) stimulate naïve T cells via major histocompatibility complex-peptide complexes on the cell surface, initiating a complex immune response process (Fig.9a) [66–68, 94, 95]. Dendritic cells are often considered the most functionally potent professional APCs in this context [67, 96]. Concurrently, naïve CD4/8+ T cells further differentiate into effector T cells with distinct biological functions under the interaction of various cytokines (Fig.9b) [97–100].

Peripheral blood (compartment B) serves as a conduit for immune cell migration between different tissues (Fig.9c) [71, 72, 83].

The tumor microenvironment (compartment C) comprises the milieu surrounding tumor cells, predominantly composed of tumor cells, immune cells, cytokines, and chemokines [101, 102]. Here, tumor cells and immune cells interact synergistically to drive disease progression [101–105]. Additionally, normal apoptosis of tumor cells, immune attack, and chemotherapy expedite the release of tumor-associated antigens (TAAs), further promoting dendritic cell activation and TAA capture [9, 10].

Bone marrow and thymus (compartment D) are regarded as primary lymphoid organs, where lymphocytes undergo further development before participating in the body’s immune function within secondary lymphoid organs [106, 107]. Together, these processes delineate the biological mechanisms of tumor immunity, as illustrated in Figure 9.

**Fig 9.**
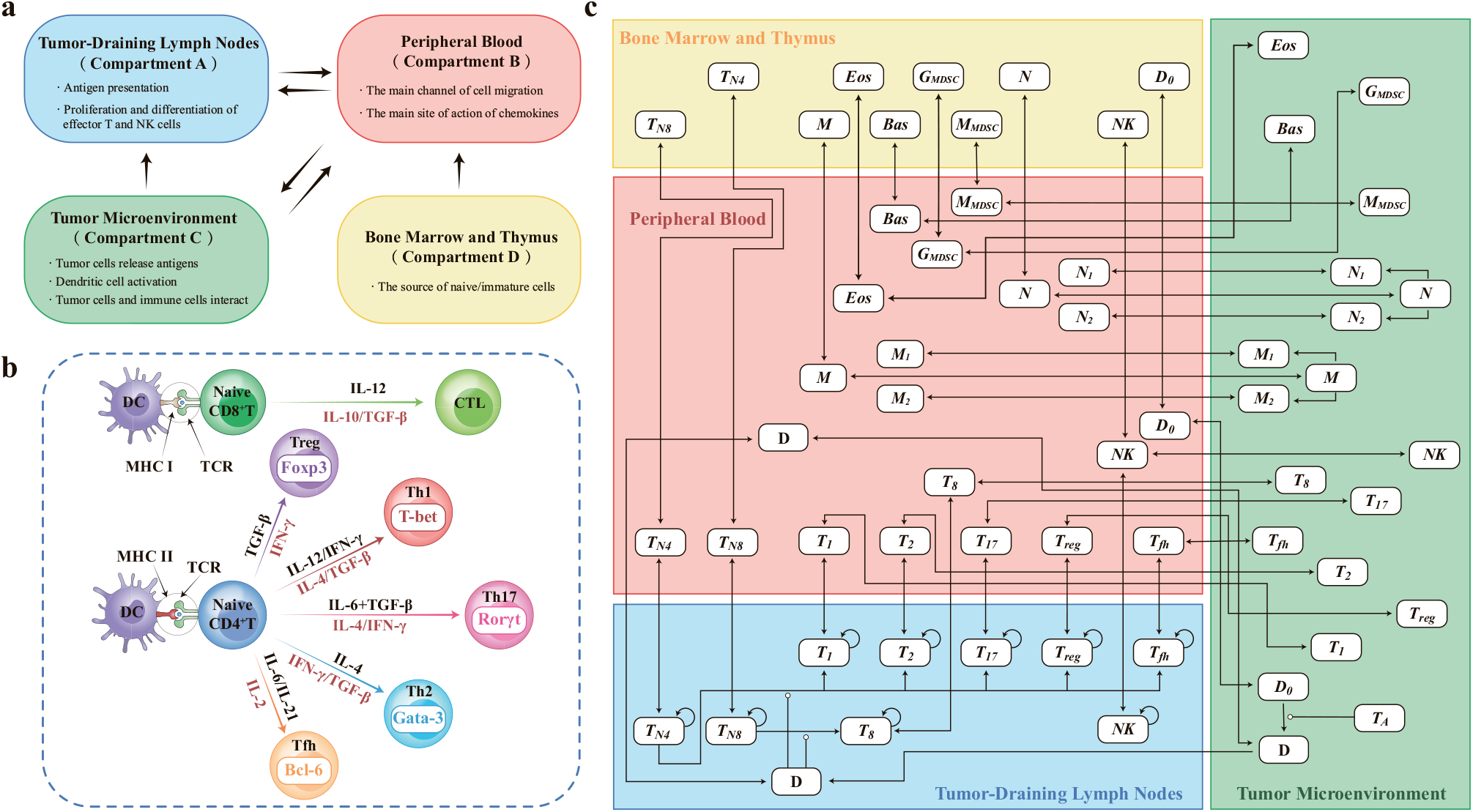
The biological mechanisms of tumor immunity. **a**. The tumor immune response comprises four compartments: tumor-draining lymph nodes (compartment A), peripheral blood (compartment B), tumor microenvironment (compartment D), and bone marrow and thymus (compartment D). Immune cells migrate between compartments via the body’s intrinsic circulatory mechanisms. Additionally, chemokines/chemokine receptors regulate the migration pattern and localization of immune cells between compartments. **b**. Illustration of immune cell differentiation. Within tumor-draining lymph nodes, tumor-associated antigens are presented by dendritic cells (antigen-presenting cells, APCs) to naïve CD4/8+T cells, which further differentiate into different effector T cells in response to various cytokines. **c**. Migration of immune cells between different compartments. *T*_*A*_, tumor-associated antigen; *D*_0_, immature dendritic cell; *D*, activated dendritic cell; *T*_*N*4_, naïve CD4+T cell; *T*_*N*8_, naïve CD8+T cell; *T*_8_, cytotoxic *T* lymphocyte cell; *T*_1_, Th1 cell; *T*_2_, Th2 cell; *T*_17_, Th17 cell; *T*_*reg*_, regulatory *T* cell; *T*_*fh*_, follicular helper *T* cell; *NK*, natural killer cell; *M*, monocyte; *M* 1, *M*1-type macrophage; *M* 1, *M*2-type macrophage; *N*, neutrophil; *N* 1, *N*1-type neutrophil; *N* 2, *N*2-type neutrophil; *M*_*MDSC*_, monocyte myeloid-derived suppressor cell; *G*_*MDSC*_, granulocyte myeloid-derived suppressor cell; *E*_*os*_, eosinophils; *B*_*as*_, basophilic.

### Pharmacokinetic and pharmacodynamic models

The pharmacokinetic (PK) model is crucial for quantitatively understanding how drugs behave within the body [108]. To capture these dynamics and assess the impact of drug therapy, we utilized a two-compartment PK model in this study, a widely adopted framework in PK research [109, 110]. Conceptually, this model divides the body into compartments based on drug transport rates, abstracting specific organs or tissues. The central compartment, characterized by rapid drug transport and elimination, contrasts with the peripheral compartment, where drug distribution occurs more slowly, necessitating drug return to the central chamber for metabolism and excretion. In our QCIC model, we designated the tumor-draining lymph nodes (compartment A) and the bone marrow and thymus (compartment D) as the peripheral compartments akin to those in a two-compartment PK model. Conversely, peripheral blood (compartment B) and the tumor microenvironment represent the central compartments. Consequently, we postulated the drug concentration in the tumor corresponds to that in the central compartment (Fig. 10a). The PK models for TAS-102 and bevacizumab are outlined in Figure 10b.

**Fig 10.**
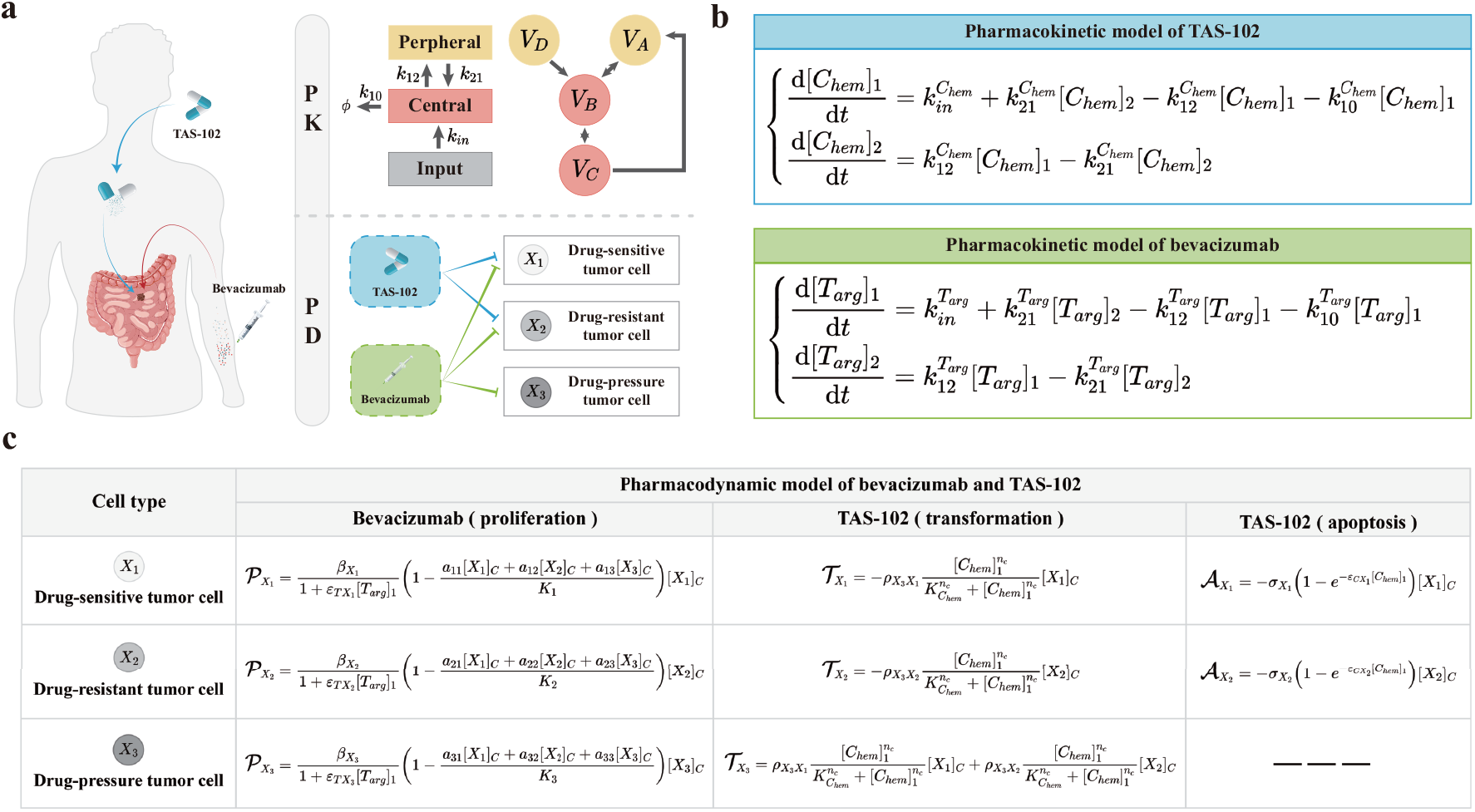
Pharmacokinetic and pharmacodynamic models. **a**. Illustration of the biological mechanisms underlying pharmacokinetic and pharmacodynamic. **b**. Pharmacokinetic model depicting the dynamics of TAS-102 and bevacizumab. **c**. Pharmacodynamic model outlining the action of bevacizumab and TAS-102. Detailed parameter explanations can be found in Supplementary Text 1 and Text 2.

The pharmacodynamic (PD) model, a mathematical tool assessing drug dosage and efficacy through drug interactions [110, 111], is integrated into our study. We hypothesized that TAS-102 primarily targets two tumor cell types: drug-sensitive tumor cells (DSTC) and drug-resistant tumor cells (DRTC). However, the emergence of drug-pressure tumor cells (DPTC), resistant to TAS-102 due to induced genetic mutations, poses a challenge (Fig. 10a). Conversely, given bevacizumab’s anti-angiogenic properties, we assumed that its principal role is to suppress tumor growth (Fig. 10a). Detailed pharmacological mechanisms for TAS-102 and bevacizumab are provided in Figure 10c.

### Virtual patient generation and in silico clinical trials

*In silico* clinical trials involve computational simulations that mimic real clinical trials, offering a method to generate virtual patient cohorts for evaluating treatment outcomes [38–50]. To generate these virtual patient cohorts, certain parameters of the QCIC model are varied while others are held constant. We identified 21 parameters across 7 categories, encompassing factors like the carrying capacity of naïve T cells, antigen activation rate, immune checkpoint expression rate, tumor proliferation rate, tumor carrying capacity, tumor apoptosis rate, and drug correlation coefficient, as key determinants of tumor heterogeneity (see Supplementary Text 2 and Text 3). Supplementary Figure 3 illustrates the results of the global sensitivity analysis conducted on these tumor heterogeneity parameters.

The values and ranges of these parameters were derived from established mathematical models and clinical trials focusing on metastatic colorectal cancer (mCRC). Further refinement was achieved by calibrating parameter distributions using data from 10 selected clinical trials [55–64] (see Supplementary text 3). The algorithm process is outlined in Figure 11.

**Fig 11.**
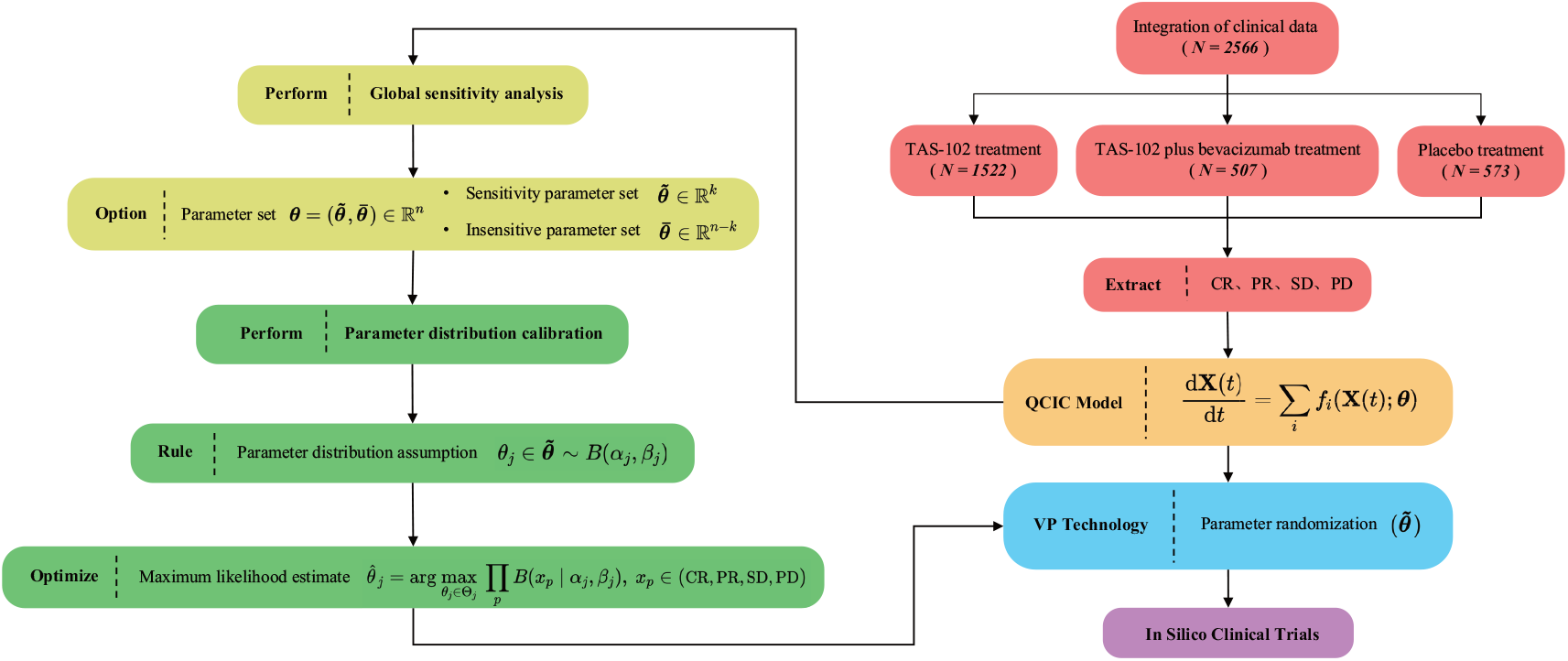
Virtual patient generation and in silico clinical trials. The process begins with obtaining clinical trial data encompassing complete response (CR), partial response (PR), stable disease (SD), and progressive disease (PD) outcomes. Subsequently, a global sensitivity analysis is conducted to identify tumor heterogeneity parameters. These parameters, assumed to follow beta distribution, are sequentially calibrated using experimental data and maximum likelihood estimation. Finally, virtual patient cohorts are generated via randomized parameter sampling for conducting *in silico* clinical trials.

### Constructing evaluation indexes for tumor clinical efficacy

Initially, we adopted the Response Evaluation Criteria in Solid Tumors (RECIST) version 1.1 as the standard for assessing tumor progression [93]. Subsequently, adhering to RECIST V1.1 guidelines, we primarily quantified six indices: complete response (CR), partial response (PR), stable disease (SD), progressive disease (PD), overall response rate (ORR), and disease control rate (DCR). Compare the calculation results with clinical data to verify the accuracy of the model.

To gauge the dynamic changes in solid tumors among virtual patients, we introduced the treatment response index (TRI) defined as:

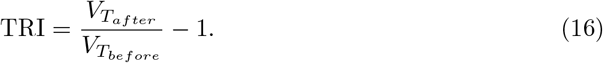

Here, *V*_*T*_ represents the tumor volume and is calcualted as:

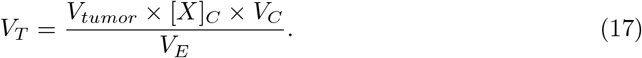

In this equation, [*X*]_*C*_ denotes tumor cell density, *V*_*C*_ stands for the volume of the tumor microenvironment (TME), *V*_*tumor*_ represents the volume of an individual tumor cell, and *V*_*E*_ indicates the tumor volume fraction. Notably, *V*_*tumor*_ = 2.572 *×* 10^*−*9^cm^3^/cell and *V*_*E*_ = 0.37 as previously determined [43, 48]. The TRI categories for classifying virtual patients are outlined in Table 2.

### Survival analysis based on the death probability function

The Kaplan-Meier (K-M) survival analysis is a standard non-parametric method in clinical medicine used to estimate survival probabilities at the population level by observing survival times. The K-M survival function is described by

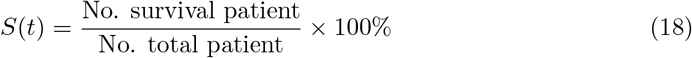

In mathematical models, the survival of virtual patients is typically evaluated by setting a threshold for either the total number of tumor cells or the tumor volume [112].

However, due to the patient variability and tumor heterogeneity, the likelihood of patient death can significantly vary. To address this inherent randomness, we introduced a death probability function (DPF):

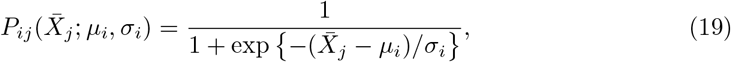

where *i* represents the *i*-th type of treatment option, *µ*_*i*_ and *σ*_*i*_ represent the shape parameters of the *i*-th type of treatment option, and 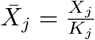 represents the ratio of the dominant tumor cell type *X*_*j*_ to the carrying capacity *K*_*j*_. *P*_*ij*_ describes the death probability caused by the dominant tumor cell type *X*_*j*_ under the *i*-th treatment option. The ranges of values for these shape parameters are detailed in Supplementary Text 4.

## Supporting information

**S1 Appendix. Supplementary materials**.

## Notes

### Competing Interest Statement

The authors have declared no competing interest.

